# Alpha-Synuclein is a Target of Fic-mediated Adenylylation/AMPylation: Implications for Parkinson’s Disease

**DOI:** 10.1101/525659

**Authors:** Anwesha Sanyal, Sayan Dutta, Aswathy Chandran, Antonius Koller, Ali Camara, Ben G. Watson, Ranjan Sengupta, Daniel Ysselstein, Paola Montenegro, Jason Cannon, Jean-Christophe Rochet, Seema Mattoo

## Abstract

During disease, cells experience various stresses that manifest as an accumulation of misfolded proteins and eventually lead to cell death. To combat this stress, cells activate a pathway called UPR (Unfolded Protein Response) that functions to maintain ER (endoplasmic reticulum) homeostasis and determines cell fate. We recently reported a hitherto unknown mechanism of regulating ER stress via a novel post-translational modification (PTM) called Fic-mediated Adenylylation/AMPylation. Specifically, we showed that the human Fic (filamentation induced by cAMP) protein, HYPE/FicD, catalyzes the addition of an AMP (adenosine monophosphate) to the ER chaperone, BiP, to alter the cell’s UPR-mediated response to misfolded proteins. Here, we report that we have now identified a second target for HYPE - alpha-Synuclein (αSyn), a presynaptic protein involved in Parkinson’s disease (PD). Aggregated αSyn has been shown to induce ER stress and elicit neurotoxicity in PD models. We show that HYPE adenylylates αSyn and reduces phenotypes associated with αSyn aggregation *in vitro*, suggesting a possible mechanism by which cells cope with αSyn toxicity.

**HIGHLIGHTS:**

- Aggregated forms of the presynaptic protein αSyn cause neurotoxicity and induce ER stress in cellular and animal models of Parkinson’s disease.
- We have identified αSyn as a novel target for the human Fic protein, HYPE, a key regulator of ER homeostasis.
- HYPE adenylylates αSyn and reduces the aggregation of recombinant αSyn
- Fic-mediated adenylylation/AMPylation is a possible mechanism by which cells cope with αSyn toxicity.

**Graphic Abstract:** 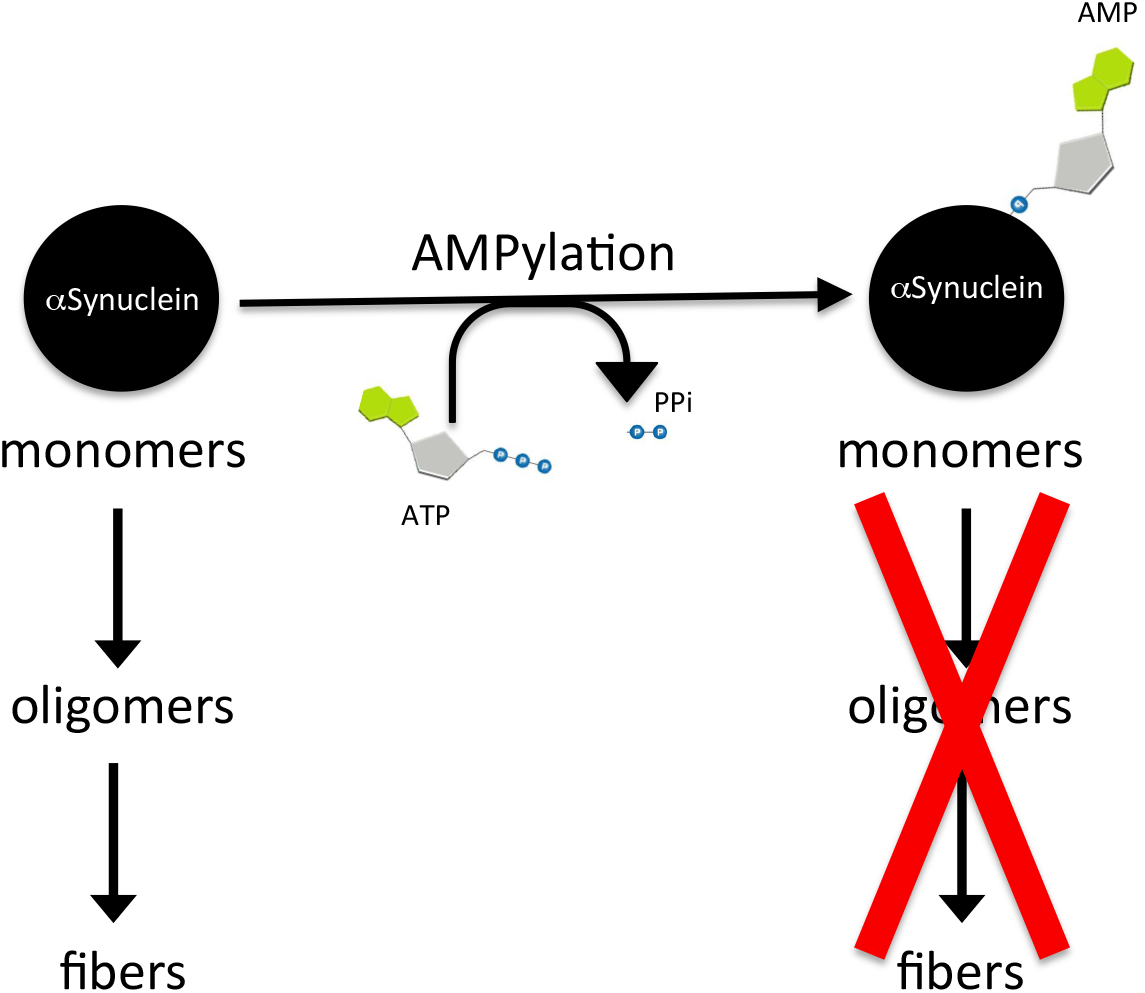

## INTRODUCTION

Post-translational modifications (PTMs) catalyzed by the Fic (filamentation induced by cAMP) family of enzymes are fast being recognized as a central regulatory paradigm governing diverse signal transduction pathways in both prokaryotes and eukaryotes [1]. Fic proteins consist of a core HxFx(D/E)(G/A)N(G/K)RxxR motif, with the Histidine being essential for catalytic activity [2]. Additionally, the activity of most Fic proteins is inhibited by inter- and intramolecular interactions with an inhibitory helix (α-inh), characterized by an (S/T)xxxE(G/N) motif [3]. We previously discovered that some bacterially secreted Fic proteins could induce host cell toxicity by covalently modifying mammalian Rho GTPases RhoA, Rac1, and Cdc42 with AMP (adenosine monophosphate) [2], [4]. This adenylylation or ‘AMPylation’ event renders the Rho GTPases inactive, thereby inducing cytoskeletal collapse and allowing the bacteria to evade phagocytosis [5], [6]. We further showed that the activity of Fic proteins was conserved in eukaryotes. Specifically, we demonstrated that the sole Fic protein in humans, HYPE (Huntingtin yeast interacting protein E) or FicD, functions as an adenylyltransferase [2, 4, 7]. HYPE is a 52kDa protein and is classified as a Class II Fic protein, consisting of a canonical Fic domain and an α-inh sequence that lies N-terminal to its Fic motif (Figure 1A). Accordingly, a mutation of HYPE’s His363 to Ala renders it enzymatically inactive, while a mutation of its Glu234 to Gly renders it constitutively active [3], [7]. We showed that HYPE localizes to the lumen of the ER (endoplasmic reticulum) and adenylylates the Hsp70 chaperone, BiP (Binding immunoglobulin protein) [7]. BiP monitors protein folding in cells and serves as a sentinel for the activation of an ER stress response pathway called the UPR (unfolded protein response)[8].

**Figure 1:**
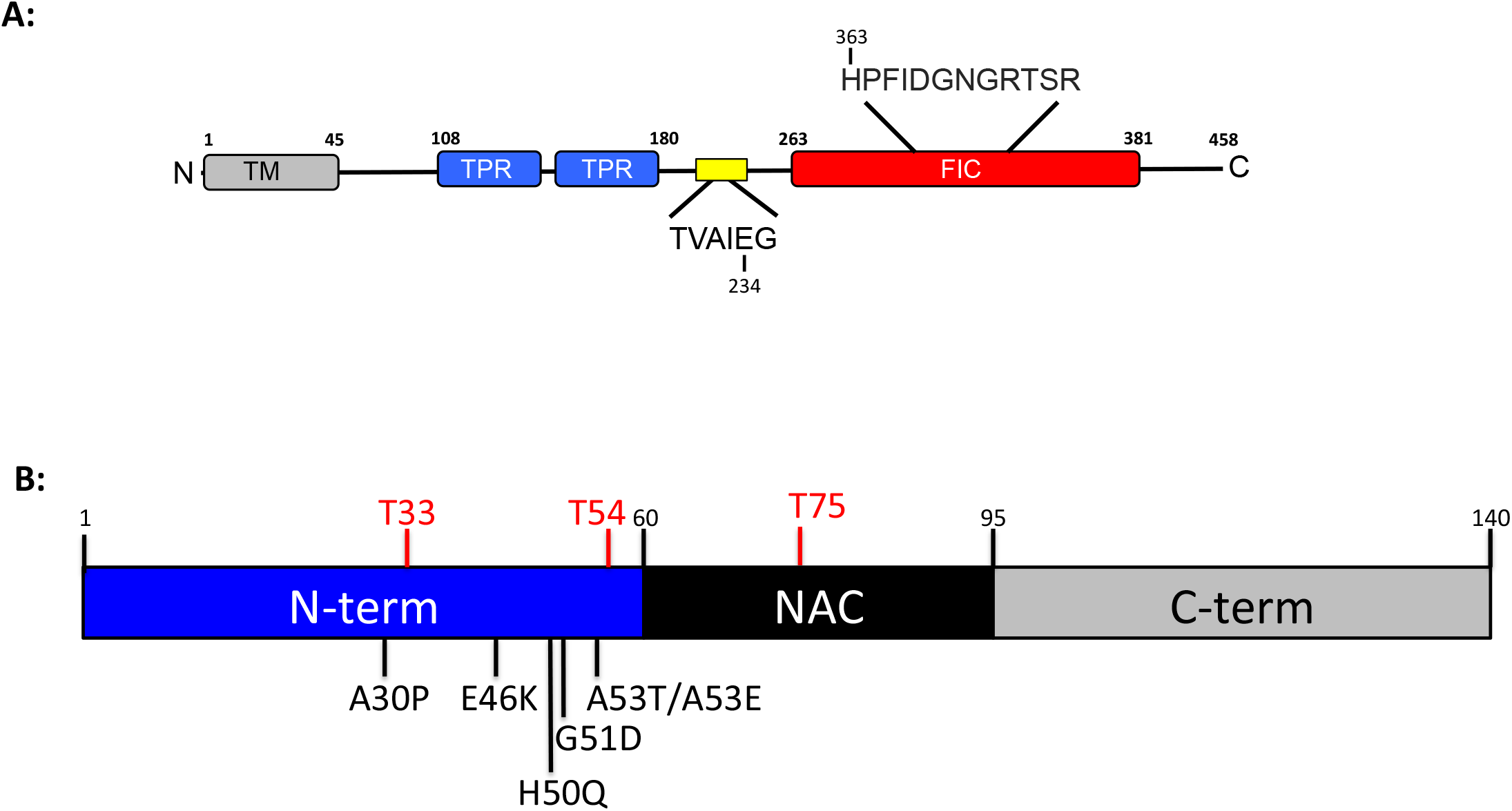
**A**) Schematic representation of human HYPE. HYPE’s Fic motif with the catalytic H363 and its α-inh domain (yellow box) with E234 are indicated. SS/TM, signal sequence/transmembrane domain. TPR, tetratricopeptide domain. Fic, filamentation induced by cAMP domain. **B**) Schematic representation of human alpha-Synuclein (αSyn). Familial mutants associated with Parkinson’s disease (A30P, E46K, H50Q, G51D, A53E, and A53T) are indicated. Sites of adenylylation (T33, T54, and T75) are indicated in red. αSyn’s N-terminal (N-term), central hydrophobic NAC (non-Abeta component of Alzheimer’s disease amyloid), and C-terminal (C-term) domains are shown. Schematic representations for HYPE and αSyn are not to scale.

During disease, cells experience various stresses that manifest as an accumulation of misfolded proteins and eventually lead to cell death [9]. To combat this stress, cells activate the UPR pathway, which initially functions to restore normalcy to the cell. However, upon sustained stress, the UPR triggers apoptosis. Thus, maintenance of ER homeostasis is a critical aspect of determining cell fate and requires a properly functioning UPR. It was recently reported that while E234G-HYPE adenylylates BiP, WT-HYPE serves to reverse HYPE-mediated adenylylation of BiP [10]. Thus, WT-HYPE is classified as a de-AMPylase and a H363A mutation in the wild type background renders an inactive de-AMPylase. Together, the adenylyltransferase and de-AMPylase activity of HYPE on BiP highlights the important role HYPE plays in tightly regulating UPR and ER homeostasis in response to protein misfolding.

Proper protein folding entails a process where the nascent polypeptides are folded into a three-dimensional functional identity. The organized native structure of the protein is at its energy minimum state while the unfolded polypeptide has higher free energy [11]. Molecular chaperones in the cytosol and ER, such as BiP, bind misfolded proteins with varying specificity and mediate their folding [12]. Any changes in the system that destabilize the folding process lead to the formation of misfolded species that lack a thermodynamically stable structure and hence have a propensity to aggregate [13]. These aggregates are prone to further aggregation by a process called nucleation, thus increasing in size at an accelerated rate [13]. Such aggregates are a hallmark of various degenerative diseases like Parkinson’s disease (PD), Alzheimer’s disease, Huntington’s disease, type II diabetes, cystic fibrosis and others [13]. Several lines of evidence suggest that accumulation of specific protein aggregates is toxic for cells [14].

PD is a progressive neurodegenerative disorder characterized by tremors, bradykinesia, rigidity and instability of posture [15]. These motor symptoms are thought to arise due to the toxic accumulation of aggregates of a presynaptic protein - alpha-Synuclein (αSyn) - in dopaminergic neurons of the *substantia nigra* in the midbrain [16]. αSyn is encoded by the SNCA gene and consists of a 140 amino acids (Figure 1B). It is an intrinsically disordered protein that consists of an amphipathic N-terminal region and an acidic C-terminal region flanking a highly hydrophobic central domain, also referred to as the NAC (Non-Amyloid-beta Component of Alzheimer’s disease) domain. Because of its natively unfolded structure, αSyn has a high propensity to aggregate[13]. Further, point mutations in the SNCA gene that target Ala53 in particular and those that result in Ala53Thr, Ala53Glu, Ala30Pro, Glu46Lys, His50Gln, and Gly51Asp substitutions, as well as duplication or triplication of the gene, lead to familial forms of PD due to enhanced aggregation of the protein [17]. Acetylation, methylation, O-GlcNAcylation, and phosphorylation of αSyn are also reported, suggesting possible roles of PTMs in PD [18] [53]. αSyn fibrils accumulate in structures called Lewy bodies in surviving neurons [19]. Although Lewy bodies are a hallmark neuropathological feature of PD, evidence suggests that the intermediate oligomeric species capable of forming fibrils may be more toxic than Lewy bodies, potentially because of their ability to permeabilize lipid membranes [20–23]. Such αSyn oligomers can also diffuse into the ER lumen and interact directly with BiP. Accumulation of αSyn oligomers results in ER stress and activation of UPR in affected neurons [24, 25].

Our discovery that HYPE adenylylates BiP established HYPE as a new player in UPR regulation in response to ER stress. We and others have now demonstrated that adenylylation alters BiP’s ATPase activity and activation of the downstream UPR cascades to reinstate ER homeostasis [7], [26–28]. Given this critical role for HYPE in how cells cope with stress from misfolded proteins and the direct correlation between αSyn accumulation, ER stress and PD progression, we reasoned that HYPE may play a role in PD, possibly via UPR regulation or direct interaction with αSyn. Indeed, a role for HYPE in neurodegeneration was recently reported in a *C. elegans* model, where activation of HYPE’s adenylyltransferase activity was shown to induce neuroprotective aggregative effects of proteins like amyloid beta (Aβ), mutant Huntingtin (m-Htt), and αSyn through its manipulation of cytosolic Hsp70 chaperones [29]. We, too, have shown previously that HYPE can directly bind to misfolded proteins [7], and global proteomic analyses of adenylylated peptides by mass spectrometry indicate targets for HYPE other than BiP and related heat shock proteins [30, 31]. However, αSyn has never been identified as a target for HYPE or known to be adenylylated.

Here, we report αSyn as a bona fide novel target for HYPE. We show that HYPE directly binds αSyn *in vitro* and adenylylates it at threonine residues predominantly in its N-terminus. Accordingly, analysis of rat midbrain sections and primary cultures reveals that HYPE is enriched in neurons of the *substantia nigra* and colocalizes with markers for dopaminergic neurons, sites where αSyn aggregates during PD. Importantly, adenylylation alters the structure of αSyn fibrils and leads to inhibitory effects on αSyn fibrillation and αSyn-mediated membrane permeabilization. Collectively, these results suggest that HYPE-mediated αSyn adenylylation may be a mechanism by which affected cells cope with αSyn toxicity. This is the first report identifying αSyn as a target for HYPE, and reinforces a role for Fic-mediated adenylylation/AMPylation in neurodegeneration. Importantly, our data shed light on a possible direct means of modifying and reducing toxicity from misfolded aggregates, which may function in parallel with HYPE’s effect on chaperones.

## RESULTS

### HYPE is enriched in the *substantia nigra* in rat midbrain sections

Global transcriptomic and proteomic studies indicate that HYPE is expressed at extremely low levels in all tissues, including neurons [2]. Indeed, endogenous HYPE is rarely detected in most epithelial cells like HEK293 or HeLa by standard Western blotting or immunofluorescence techniques [7]. To assess HYPE expression in neurons, we performed immunohistochemistry on sections of rat midbrain using antibody generated against bacterially purified full length human HYPE, where human and rat HYPE (FicD) share 90% amino acid sequence identity. We observed strong staining for HYPE in sections corresponding to the *substantia nigra*, a region where neuronal toxicity occurs due to αSyn aggregation during PD (Figure 2A). HYPE’s staining pattern within the *substantia nigra* appeared neuronal under higher magnification analysis (Figure 2A, red and blue boxes, and supplemental Figure 1). We also saw evidence of HYPE staining in the pyramidal neurons of the hippocampus (as indicated in Figure 2A).

**Figure 2:**
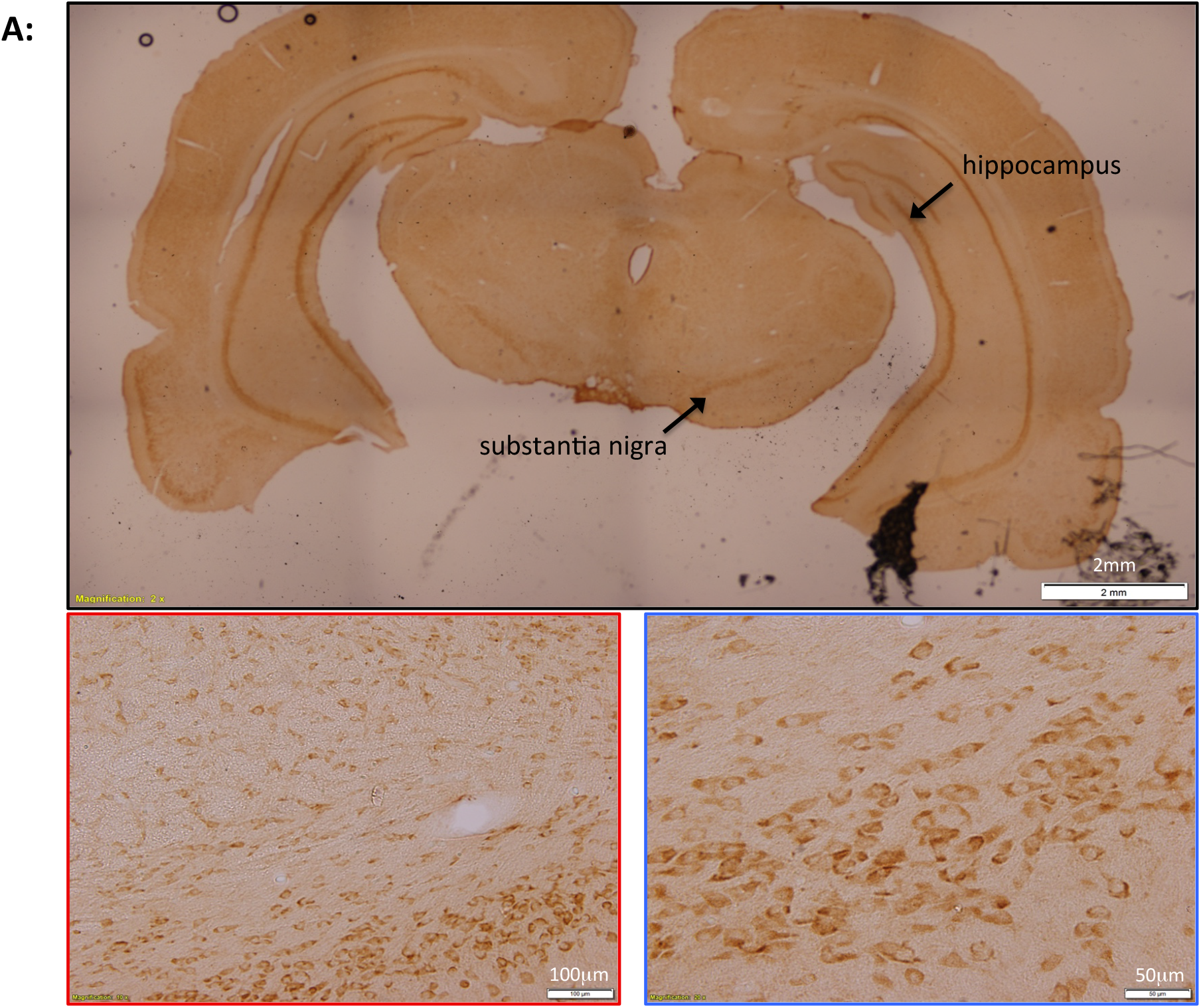
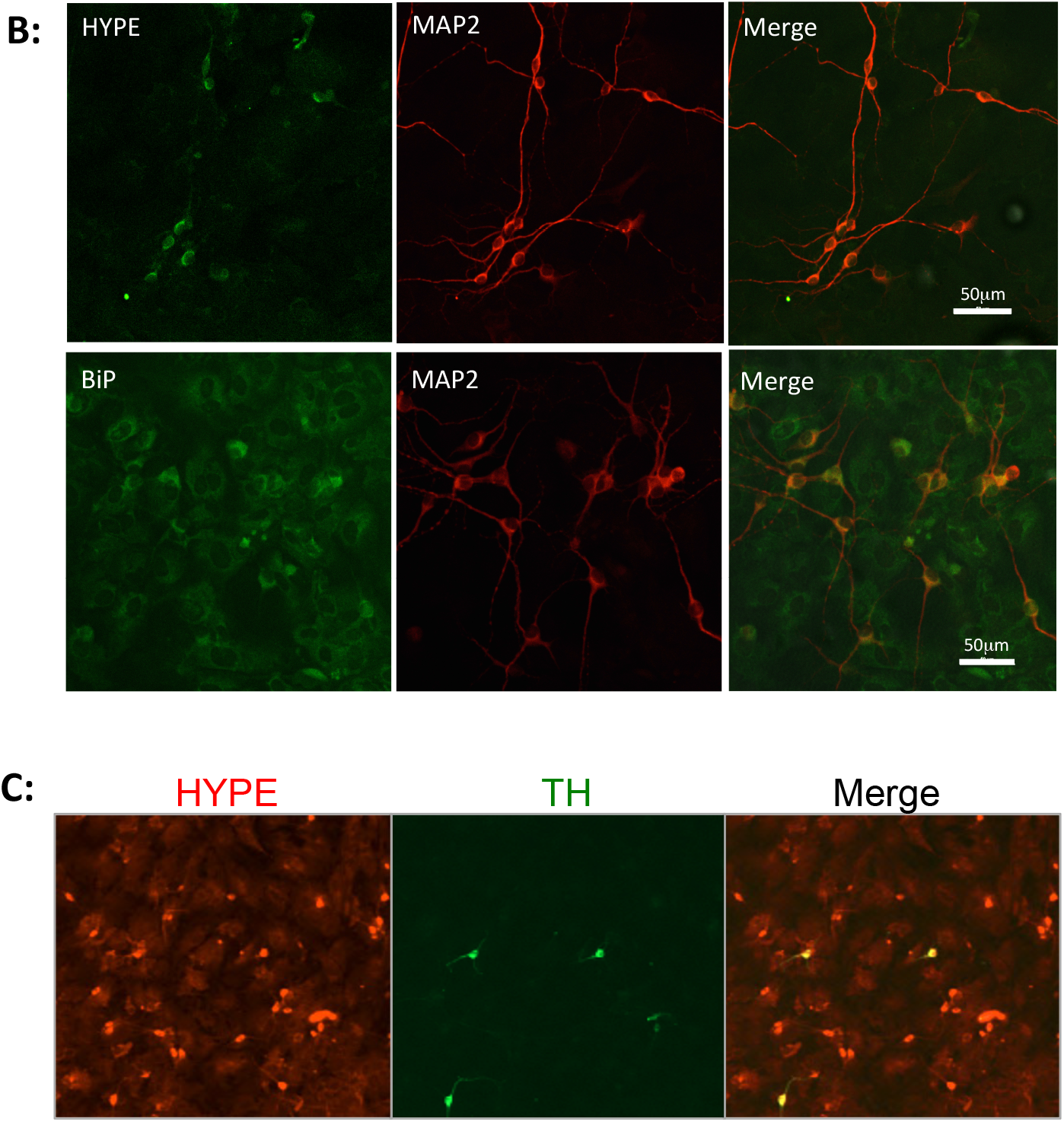
HYPE is expressed in adult and embryonic rat midbrain neurons. **A**) Top panel: Coronal section of a rat brain probed with antibody to HYPE and visualized with peroxide-based secondary staining shows strong HYPE expression in the *substantia nigra* and hippocampus. Scale bar = 2 mm. Bottom panels: Higher magnification views of HYPE staining in the nigral region. Scale bars in the red and blue boxes represent 100 μm and 50 μm, respectively. **B**) Immunofluorescence imaging of primary midbrain cultures from E17 rat embryos shows colocalization of HYPE (green, top panel) or another ER lumenal protein, BiP (green, bottom panel), with the neuronal marker MAP2 (red). HYPE shows a preferential neuronal expression, whereas BiP is expressed highly in both neurons and surrounding glial cells. Both HYPE and BiP selectively localize to the cell body in neurons. Scale bar = 50 μm. **C**) Immunofluorescence showing co-localization of HYPE (red) with the dopaminergic neuronal marker TH (tyrosine hydroxylase; green).

### HYPE is expressed in nigral dopaminergic neurons and localizes to the cell body

Toxicity induced by αSyn aggregation predominantly affects nigral dopaminergic neurons [15]. We, therefore, assessed the localization of HYPE in primary dissociated neurons derived from embryonic rat midbrain. Immunofluorescence analyses of HYPE revealed that it colocalizes with the neuronal marker MAP2 (microtubule-associated protein 2) as detected by fluorescence microscopy, confirming HYPE expression in neurons (Figure 2B), whereas HYPE expression in surrounding non-neuronal cells (consisting primarily of astrocytes) was much weaker. Interestingly, compared to MAP2, which is expressed across both the cell body and in neurites, HYPE is detected predominantly in the neuronal cell body, but not to a large extent in the dendrites (Figure 2B, upper panels). In contrast to HYPE, the chaperone protein BiP (also an ER lumenal protein) displays ER-specific expression in all parts of the neuron and is present at higher levels in surrounding astrocytes (Figure 2B, lower panels and Supplemental Figure 2). Finally, we observed HYPE expression in dopaminergic neurons that stained positive for tyrosine hydroxylase (TH) (Figure 2C), indicating that HYPE is expressed in neurons that are most vulnerable in PD. Together, these data support our hypothesized link between HYPE activity and its relevance to PD.

### HYPE interacts with αSyn and adenylylates it *in vitro*

αSyn is known to localize to the ER lumen and interact with BiP [24, 32, 33]. We previously reported that HYPE interacts with and adenylylates BiP in the ER lumen, and alters UPR activation presumably by altering BiP’s ability to refold misfolded proteins [7]. Additionally, we found that HYPE itself has the ability to bind to misfolded proteins [7]. Thus, αSyn is correctly situated in the ER to allow interactions with HYPE. Given HYPE’s role in UPR regulation and its expression in regions of the brain that are associated with aggregation of misfolded αSyn, we asked whether HYPE directly interacts with αSyn. Complementary co-immunoprecipitation experiments using SH-SY5Y neuroblastoma-derived cells with antibody to αSyn or to HYPE revealed a direct interaction between the two proteins (data not shown). Our experience with HYPE informs us that in its enzymatically active state, it interacts transiently with its substrates (like BiP); however, transient interactions can be captured *in vitro* by assessing interactions using the catalytically dead E234G/H363A-HYPE mutant, which functions as a substrate trap [7]. We, therefore, conducted biolayer interferometry with bacterially expressed and purified His-tagged αSyn and untagged E234G/H363A-HYPE, with αSyn immobilized on the anti-His antibody sensor, to determine HYPE’s binding efficiency for αSyn. E234G/H363A-HYPE displayed sustained binding to αSyn with a Kd of 15.3 + 0.04μM (Figure 3A).

**Figure 3:**
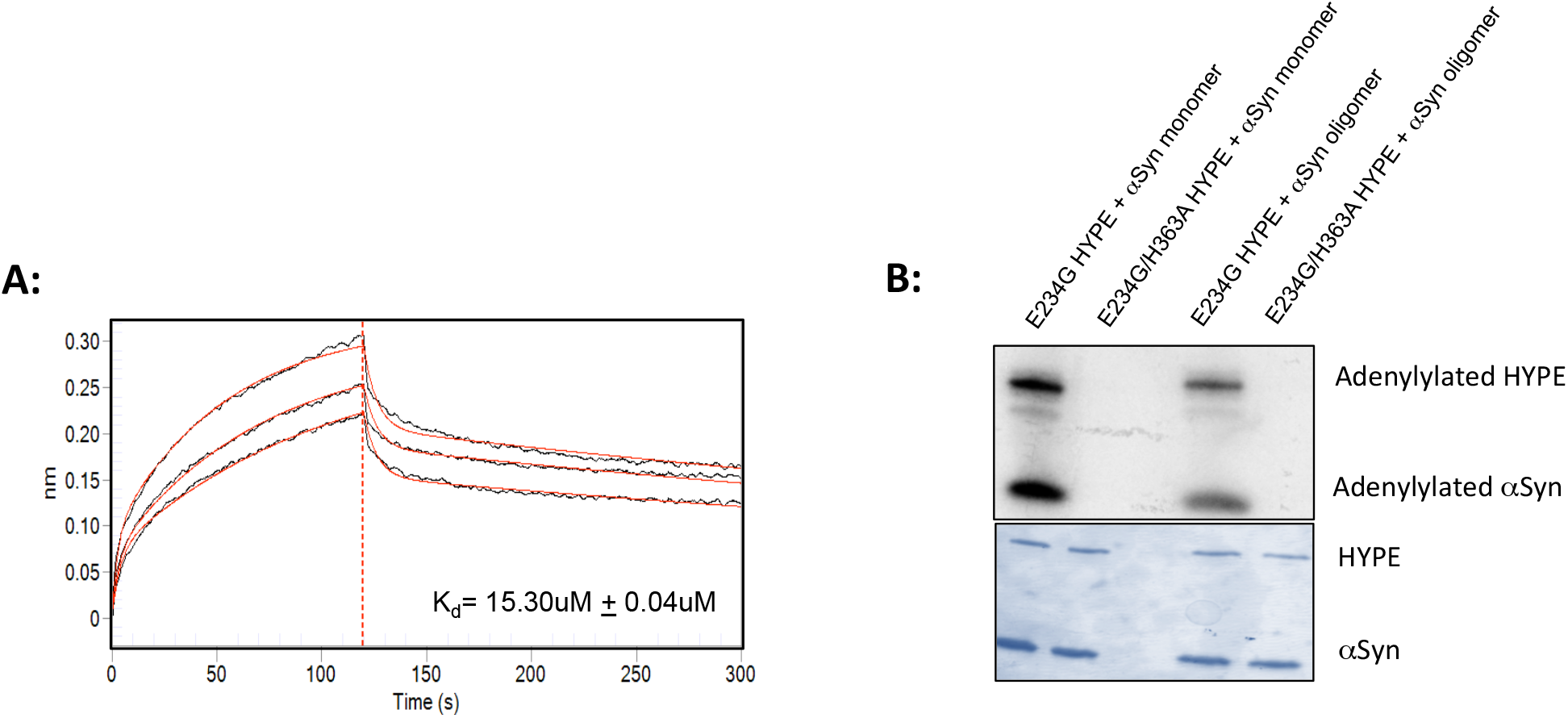
HYPE binds to and adenylylates αSyn. **A**) Steady state binding affinity determination for E234G/H363A-HYPE incubated with His6-tagged αSyn immobilized on HIS1K sensors were obtained by Biolayer Interferometry. **B**) *In vitro* adenylylation assay with purified HYPE and αSyn using α^32^P-ATP as a nucleotide source showing adenylylation of αSyn by enzymatically active E234G-HYPE but not by its catalytically dead mutant E234G/H363A-HYPE.

We next asked if HYPE could adenylylate αSyn. For this, untagged αSyn was purified as oligomeric and as low molecular weight, mostly monomeric species (by passing the αSyn solution prepared by dissolving the lyophilized protein over a 100kD-cut off filter). Monomeric and oligomeric αSyn were incubated with E234G-HYPE or its enzymatically inactive E234G/H363A mutant in an *in vitro* adenylylation reaction using α-^32^P-ATP as a nucleotide source. Both monomeric and oligomeric forms of αSyn were efficiently adenylylated by enzymatically active E234G-HYPE but not by E234G/H363A HYPE (Figure 3B). Even though equal concentrations of monomeric and oligomeric αSyn were used in this reaction, monomeric αSyn appeared to be more efficiently adenylylated, as determined by stronger intensity of radiolabelled bands corresponding to αSyn on the autoradiogram (Figure 3B). These data are the first indication of αSyn as a novel target for HYPE-mediated adenylylation.

### HYPE adenylylates αSyn at threonine residues predominantly in its N-terminus

Monomeric αSyn incubated with E234G-HYPE in an *in vitro* adenylylation reaction as described above was digested with trypsin or chymotrypsin and subjected to LC-MS/MS mass spectrometric analysis (see Methods). Peptides with a mass shift corresponding to the addition of an AMP were found to be modified at amino acids T33, T54, and T75. Representative spectra for peptides with an AMP modification at T33, T54, and T75 showing peaks at m/z 348.08 representing the protonated form of AMP and neutral losses from the peptides of adenine, adenosine or AMP are shown in Figures 4A, B, and C. Interestingly, nearly all the αSyn adenylylation sites identified lie in the N-terminus domain, a region critical for interacting with the membrane and needed for stabilizing αSyn [34].

**Figure 4:**
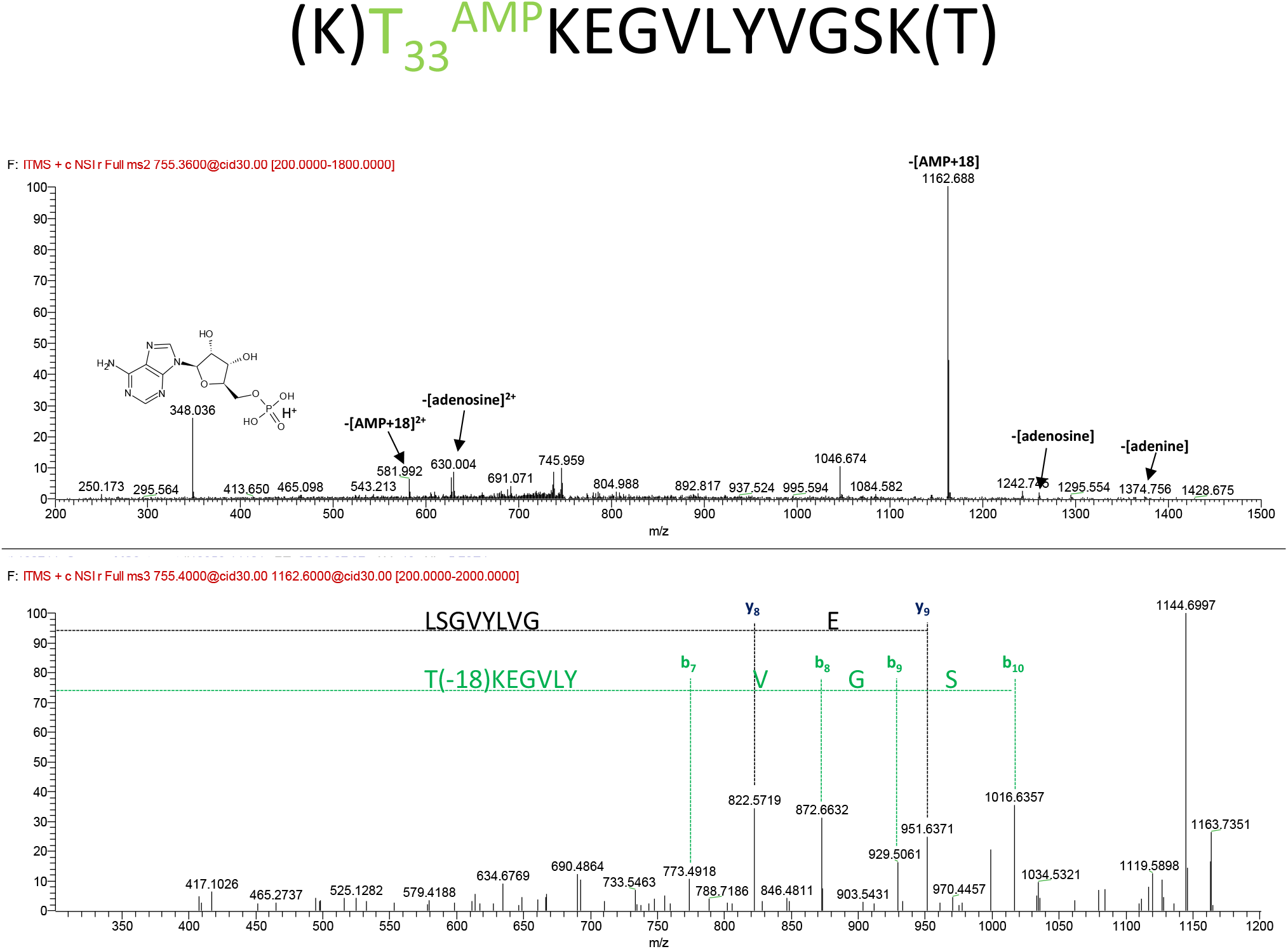
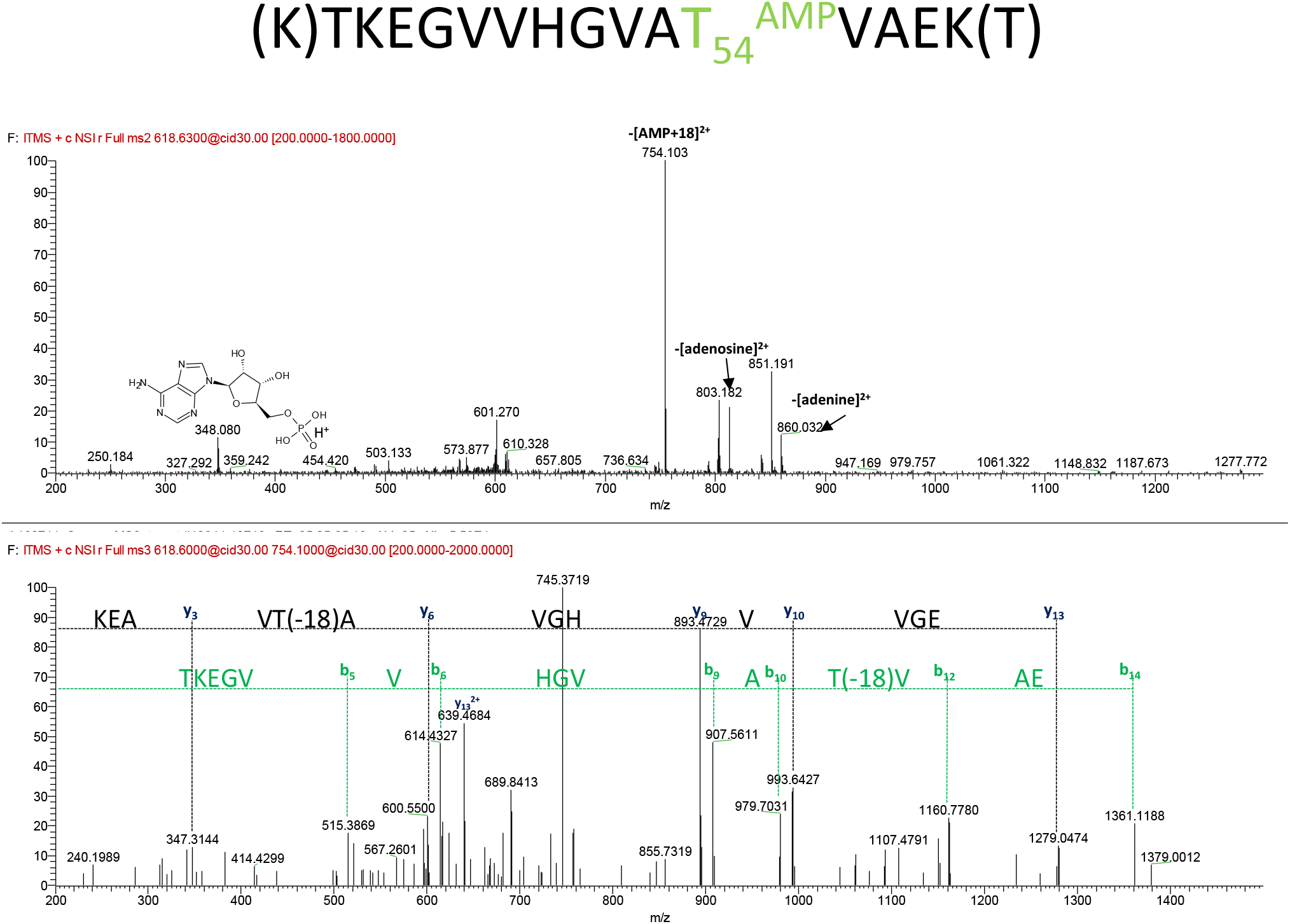
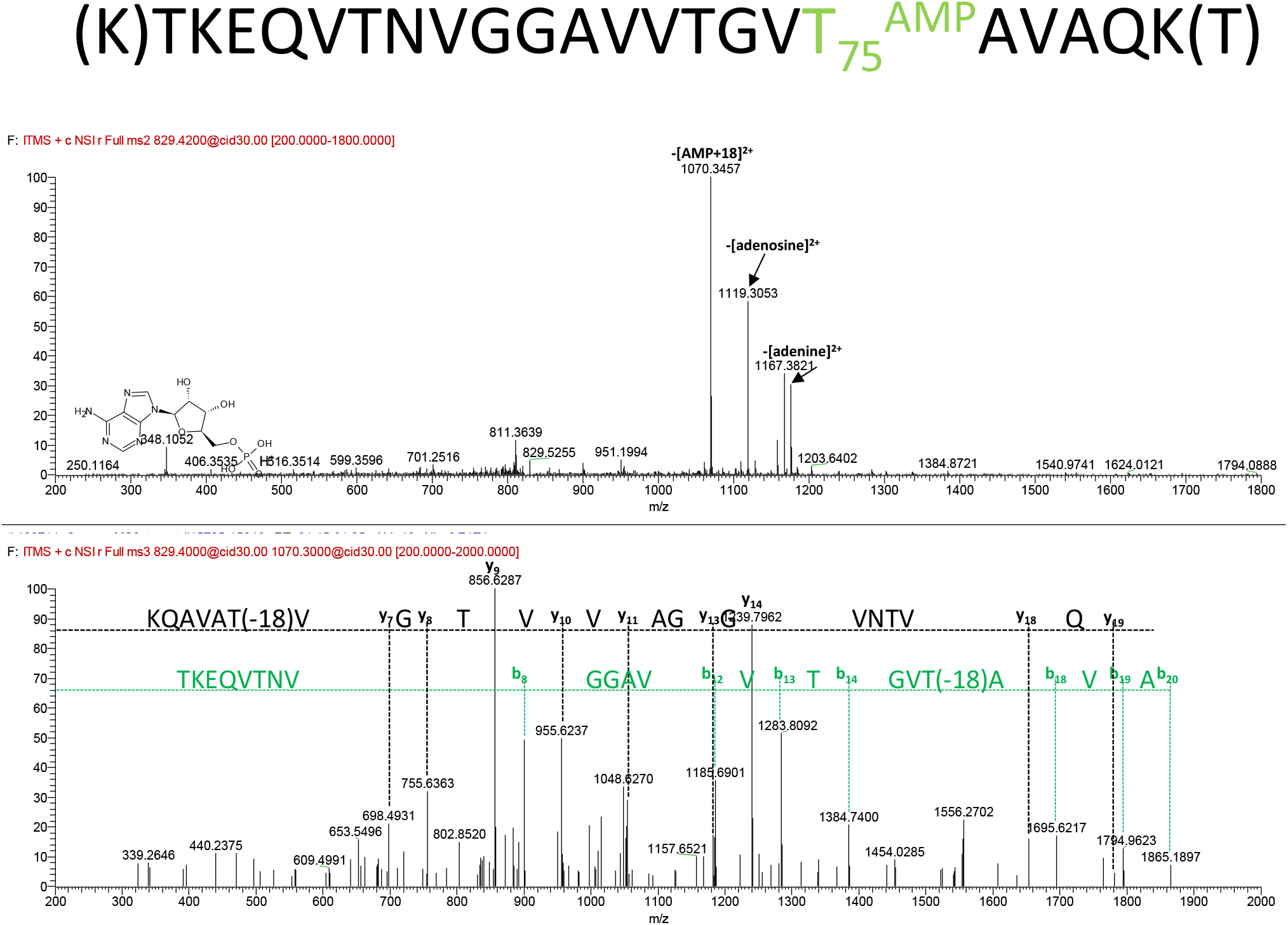
Fragmentation of AMP-modified peptides. Figures A, B, and C represent the fragmentation of AMP-modified peptides in MS2 (top) and MS3 (bottom) of the neutral loss fragment of AMP. A) MS2 and MS3 fragmentation of peptide T^33^AMPKEGVLYVGSK. B) MS2 and MS3 fragmentation of peptide TKEGVVHGVAT^54^AMPVAEK. C) MS2 fragmentation of m/z 755.36 of peptide TKEQVTNVGGAVVTGVT^75^AMPAVAQK. Annotated are fragments that lose AMP, adenosine or adenine, either from 1+ ions or 2+ ions (2+). The most abundant ion (-AMP) was subsequently isolated and fragmented again to produce fragments to identify the peptide. B and y ions are annotated as well as their respective sequences/amino acids.

### Adenylylation of αSyn shows inhibitory effects on aggregation and fibril formation

As mentioned earlier, αSyn aggregation is thought to play a key role in neurodegeneration associated with PD. Therefore, to determine the functional consequence of αSyn adenylylation, we asked whether adenylylation affects the kinetics of αSyn fibril formation. To monitor fibrillation kinetics, we conducted an *in vitro* αSyn fibrillation assay using thioflavin T (ThT), a fluorescent dye that forms a complex with the aggregating αSyn protein. ThT fluorescence intensity directly correlates with the extent of fibrillation. Mouse WT αSyn and a human A53T αSyn clinical mutant, which are known to display an increased rate of fibrillation, were used as positive controls [35]. Data from representative fibrillation assays are presented in Figure 5A and in supplemental Figure 3. αSyn alone or first adenylylated by E234G-HYPE was allowed to incubate with ThT under shaking conditions to induce the formation of fibrils and monitored for 5 days. As expected, untreated WT-αSyn formed aggregates reaching saturation levels, as evidenced by a sigmoidal fluorescence trace with a distinct lag phase and plateau (Figure 5A, blue line). In contrast, data from five of the seven biological replicates showed that fibril formation by αSyn adenylylated by E234G-HYPE was significantly compromised, displaying markedly reduced ThT fluorescence levels (Figure 5A; compare blue line vs green line). αSyn left unmodified by incubating with catalytically inactive E234G/H363A-HYPE did not show impaired fibrillation, indicating that the mere presence of HYPE does not affect the fibrillation process (Figure 5A; purple line). Interestingly, incubation of αSyn with WT-HYPE also displayed a slight reduction in fibrillation levels in some biological replicates (Figure 5A, red line and supplementary Figure 3). Quantification of the end-point fibrillation levels, however, indicated that αSyn fibrillation levels were significantly reduced only upon incubation with the adenylylation-competent E234G-HYPE (Figure 5B).

**Figure 5:**
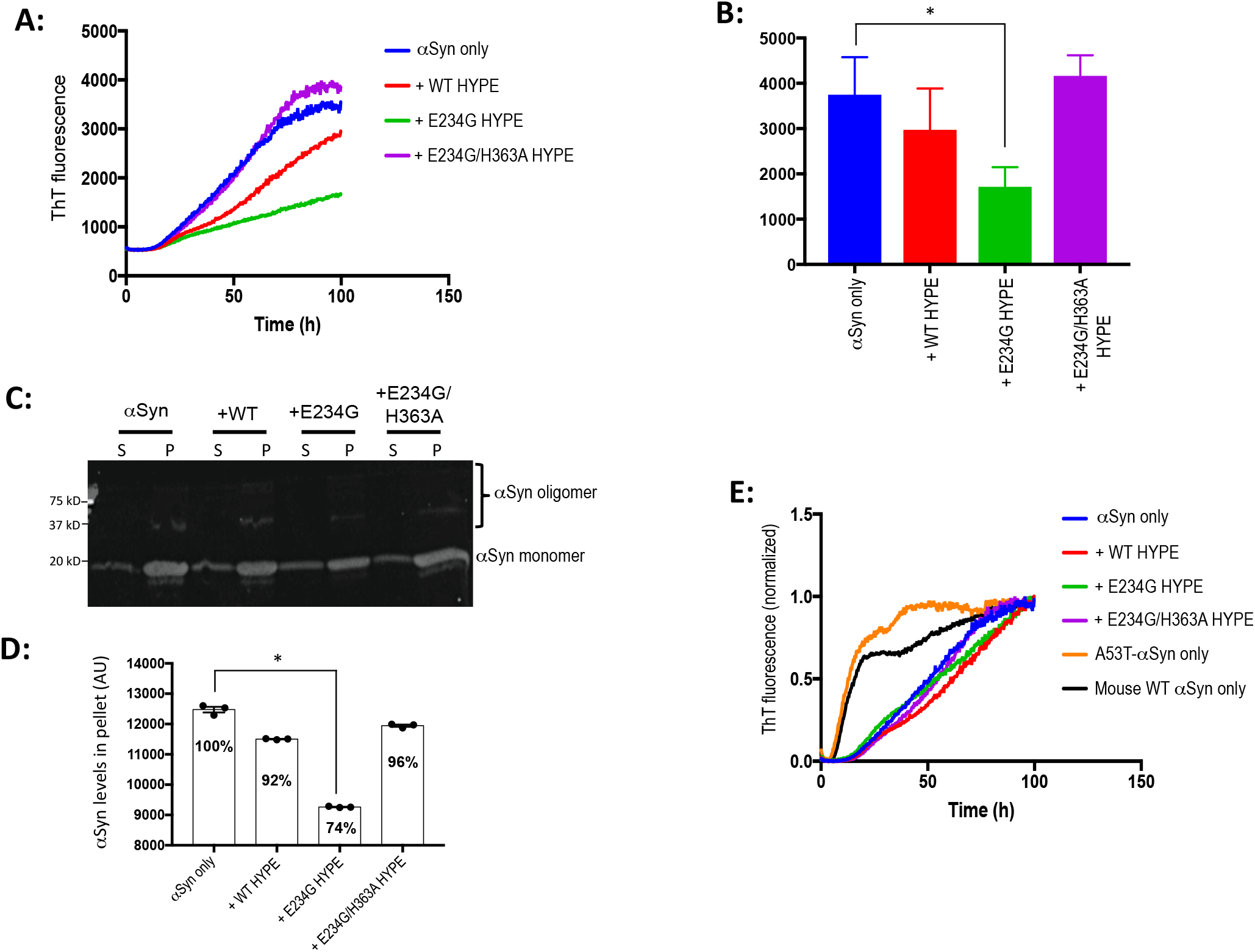
HYPE-mediated adenylylation of αSyn apparently inhibits fibril elongation without altering the lag time. **A**) Representative kinetic traces of fibrillation assays conducted with purified αSyn adenylylated in the presence of E234G HYPE (green line) or left unmodified as αSyn alone (blue line) show a significant reduction in the yield of ThT fluorescence – potentially reflecting inhibition of fibril elongation – as a result of αSyn adenylylation. Control samples consist of αSyn incubated in the presence of WT HYPE (red line) or catalytically dead E234G/H363A HYPE (purple line) prior to fibrillation. **B**) Quantification of endpoint ThT signal intensities obtained from N=3 fibrillation assays. Asterisk * indicates a p value = 0.02. **C**) Endpoint fibril samples from (A) were ultracentrifuged, and soluble αSyn in the supernatant (S) fraction and insoluble/aggregated αSyn in the pellet (P) fraction were analyzed by SDS-PAGE. **D**) Quantification of the bands corresponding to monomeric αSyn in the pellet fractions of the gel shown in (C). Adenylylation of αSyn following incubation with E234G-HYPE results in a decrease (74% vs. 100%) in the amount of αSyn seen in the pellet fraction, suggesting that more αSyn is soluble/not aggregated. **E**) Normalized ThT values indicate that the lag time of fibrillation is unaltered by HYPE-mediated adenylylation, as the curves for WT αSyn and αSyn incubated with various HYPE samples overlap. In contrast, control samples of mouse WT αSyn (dark blue line) or human A53T αSyn mutant (orange line) exhibit a shorter lag time, indicating a much faster rate of nucleation.

The fibrillation results were verified by separating and quantifying the insoluble pellet (aggregated αSyn fibrils) and supernatant (soluble, monomeric αSyn) fractions from each of the fibrillation samples. Specifically, equal amounts of end-point fibrillation samples were subjected to ultracentrifugation. The supernatant (S) and pellet (P) fractions were separated by SDS-PAGE and analyzed by western blotting with antibody to αSyn, to distinguish the monomeric and oligomeric αSyn proteins (Figure 5C). Quantification of monomeric αSyn in each of these fractions revealed that adenylylation significantly reduced the amount of insoluble αSyn (Figure 5D). Depicted in Fig 5D is the quantification of monomeric αSyn in “Pellet” fractions normalized against the corresponding αSyn only (100%) sample. The histogram corresponding to adenylylated αSyn shows decreased insolubility of 74%. Interestingly, assessment of the fibrillation data shown in Figure 5A normalized against the maximum fluorescence for each individual sample revealed a similar lag time for all samples incubated with HYPE when compared to αSyn alone (Figure 5E), whereas the lag times observed for the A53T and mouse αSyn positive controls were markedly shorter.

Taken together, these data suggest that adenylylation specifically limits the rate of elongation and the amount of αSyn fibrils formed, whereas it does not alter the lag time, a measure of the rate of nucleation.

### Adenylylation alters αSyn fibril morphology

Next, we examined the end-point fibrillation samples shown in Figure 5A by transmission electron microscopy (TEM) to confirm that amyloid-like fibrils were formed by WT and HYPE-treated αSyn, and to assess whether adenylylation had any effect on αSyn fibril structure. All the samples displayed an unbranched fibril morphology (Figure 6). In agreement with the literature [36], WT-αSyn fibrils were most frequently arranged as two fibrils (each with a diameter of ~10-15 nm) wound around each other via a helical twist characterized by a well-defined pitch length (Figures 6A). Fibrils with a similar twisted morphology were also observed in samples of αSyn incubated with adenylylation-incompetent WT-HYPE or E234G/H363-HYPE (Figures 6B and 6C, red arrows). In contrast, αSyn adenylylated by E234G-HYPE displayed a mixed population of fibrils: whereas some paired fibrils had a twisted morphology (red arrows), others were arranged in a parallel fashion without any twist (yellow arrows) (Figure 6D). These data suggest that αSyn adenylylation can interfere with the protein’s ability to form typical twisted fibrils under the conditions used here, possibly due to steric hindrance from the adenylylation PTM.

**Figure 6:**
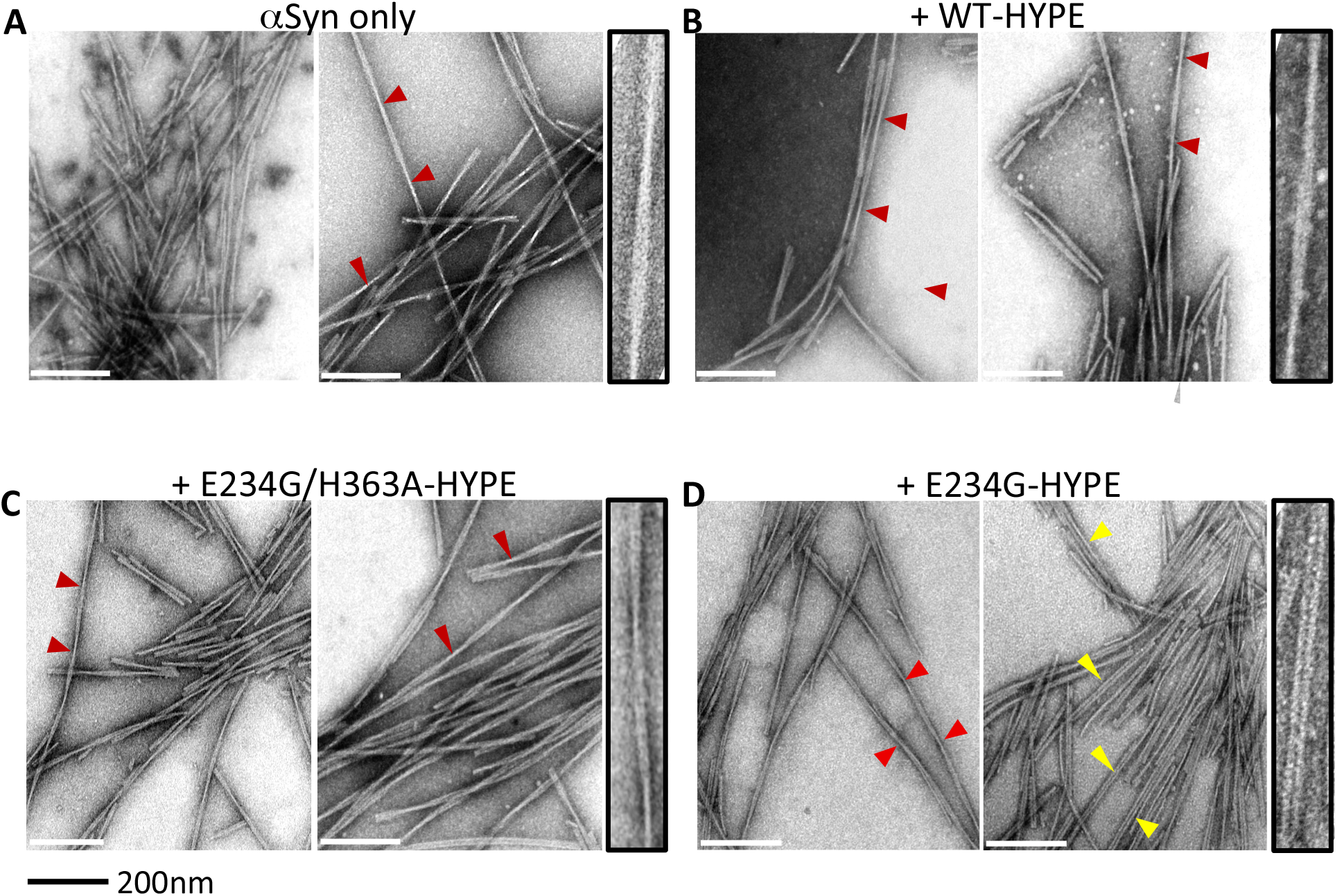
Adenylylation alters the morphology of αSyn fibrils. Transmission electron microscopy images of fibrils formed by human WT αSyn. The protein was untreated (A) or incubated with WT HYPE (B), its catalytically inactive E234G/H363A mutant, (C) or the constitutively adenylylation competent E234G mutant (D). Fibrils formed by αSyn largely consist of bundles of fibril pairs that are intertwined, with each twist occurring at regular pitch lengths (red arrows). However, significant levels of fibrils formed by adenylylated αSyn (incubated with E234G) do not display this twist and are instead structured as parallel fibril pairs (yellow arrows). Magnified images of fibrils showing one representative twist (A-C) or one parallel fibril polymorph (D) are shown beside each image. Scale 200nm.

### Adenylylation reduces membrane permeabilization

The pathological mechanism of action and physiological role of αSyn are poorly understood. Interaction with membrane surfaces has been shown to promote the process of self-aggregation and lead to disruption of bilayer integrity by lipid extraction, phenomena which could play a central role in αSyn neurotoxicity [34, 37]. We have also observed that familial mutants of αSyn that show greater neurotoxicity in primary neuronal cell cultures (e.g. A30P and G51D) also display greater vesicle disruption [34, 37]. Thus, we asked whether adenylylation alters αSyn’s ability to permeabilize the membrane. To this end, we compared unmodified WT αSyn to that incubated with WT-, E234G-, or E234G/H363A-HYPE for their ability to disrupt small unilamellar vesicles (SUV) using a calcein dye release assay. SUVs prepared from a 50:50 mixture (mol/mol) of egg phosphatidylglycerol (PG) and egg phosphatidylcholine (PC) were loaded with calcein at a self-quenching concentration and incubated with the adenylylated or unmodified αSyn variants. Permeabilization was assessed by real time measurement of the increase in fluorescence intensity caused by dequenching of the released dye into the surrounding buffer. Interestingly, αSyn adenylylated by E234G-HYPE showed a significant decrease in vesicle leakage compared to the non-adenylylated samples (Figure 7), suggesting that adenylylation may be a protective mechanism to combat the toxic effects of vesicle disruption associated with membrane-induced αSyn aggregation.

**Figure 7:**
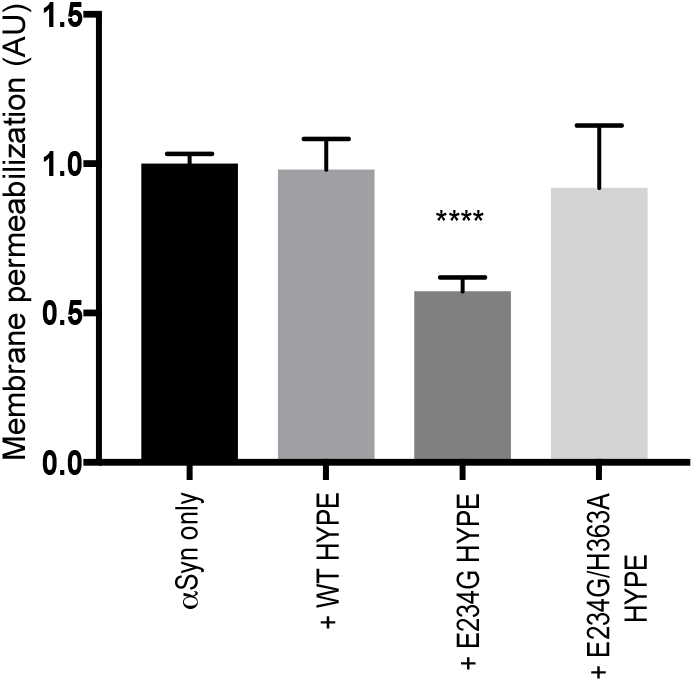
Adenylylation reduces αSyn’s ability to disrupt membranes. Calcein-loaded PG: PC (1:1) SUVs were incubated with adenylylated or non-adenylylated (control) αSyn and examined for an increase in calcein fluorescence (excitation and emission wavelengths, 485 nm and 515 nm). The data are presented as relative membrane permeabilization after 12 h, normalized with respect to non-adenylylated WT αSyn control. Mean ± SEM, n = 4 (each with 3 technical replicates).

## DISCUSSION

The discovery of Fic proteins as evolutionarily conserved adenylyltransferases has brought about a renewed interest in adenylylation/AMPylation, and underscores the importance of this previously underappreciated post-translational modification (PTM). Documentation of adenylylation in signal transduction is growing. Fic proteins represent one of three known types of adenylyltransferases, the other two being bacterial glutamine synthase adenylyltransferase (GS-AT) and the newly discovered Sel-O pseudokinase [38, 39]. PTMs catalyzed by Fic proteins are now being recognized as a universal signaling paradigm that regulates critical processes like bacterial pathogenesis, cellular trafficking, drug tolerance, and prokaryotic and eukaryotic stress response [1]. Further, our discovery of BiP as the elusive target for the human Fic protein, HYPE, has unveiled Fic-mediated adenylylation as a critical step in UPR regulation and the cell’s ability to cope with misfolded proteins [7]. Further, through its ability to manipulate chaperone activity, Fic-mediated adenylylation has recently been linked to neurodegeneration [29].

As mentioned earlier, HYPE localizes to the ER lumen. To date, BiP is the only known ER-localized physiological target for HYPE. However, some cellular and proteomic analyses have identified additional targets – such as histones and cytosolic Hsp70 and Hsp40 family chaperones - that localize to cellular compartments other than the ER [30], [31], [40]. How HYPE interacts physiologically with these non-ER localized proteins is unclear [54], though a recent report reveals that a mass exodus of ER resident proteins occurs in response to ER Ca^2+^ depletion [41]. Additionally in *C. elegans*, HYPE is reported to be partially present in the cytosol in addition to the ER lumen [29].

Here, we identify αSyn, a presynaptic protein involved in Parkinson’s disease pathogenesis, as a novel target for the human Fic protein, HYPE, and propose a direct role for adenylylation in modulating neurotoxicity. During PD, accumulation and aggregation of misfolded αSyn manifests as the formation of Lewy bodies and is thought to lead to the death of dopaminergic neurons in the *substantia nigra* region of the brain, potentially via a mechanism involving vesicle disruption. Accordingly, we see that HYPE is present in dopaminergic neurons and enriched in the *substantia nigra* of normal rat brains (Figure 2). While αSyn is predominantly a cytosolic protein, it is known to transgress membranes upon oligomerization [24, 37]. αSyn has been reported to enter the ER lumen, where it interacts with BiP and is suggested to trigger a UPR response [24]. Thus, we predict that HYPE and αSyn likely interact directly in the ER lumen. Alternatively, released cytosolic pools of HYPE may interact with αSyn. Indeed, we confirmed a direct interaction between HYPE and αSyn using immunoprecipitation and biolayer interferometry. While this binding affinity was relatively weak (in the μM range; Figure 3), it is in keeping with the binding affinity of HYPE for BiP [52]. It remains to be determined whether in a physiological setting, the interaction between HYPE and αSyn could be enhanced by the presence of BiP.

Our LC-MS/MS analysis identified three sites of adenylylation on αSyn – T33, T54 and T75. We attempted to validate these sites *in vitro* using site directed mutants of each of these sites. However, these adenylylation site-mutants of αSyn could still be modified in an *in vitro* adenylylation reaction (data not shown), likely because of secondary-site modifications, as is often seen with Fic and other enzymes [6] [7]. Interestingly, all the adenylylation sites on αSyn that we identified localized to domains of αSyn that are critical for membrane association and have also been reported as sites of O-GlcAcylation [53]. Primarily an intrinsically disordered protein, binding to the phospholipid membrane induces a structural rearrangement causing αSyn to form a helical structure [42–45]. A partial disruption of αSyn-membrane interactions contributes to αSyn aggregation at the membrane surface, a process that is apparently coupled to vesicle disruption [34, 37, 42, 46, 47]. We, therefore, assessed the physiological implications of αSyn adenylylation in PD using an *in vitro* liposome-based assay for measuring phospholipid membrane permeability, and found that adenylylation of αSyn reduces αSyn-mediated membrane permeabilization (Figure 7). In a previous study, substitutions at threonine residues in the amphipathic helical domain, including T33, were found to disrupt αSyn membrane binding [48]. By analogy with these earlier findings, adenylylation of αSyn threonine residues could perturb αSyn-lipid interactions. This effect, as well as a potential adenylylation-mediated disruption of interactions between neighboring αSyn molecules at the membrane surface, could lead to a decrease in the protein’s ability to elicit vesicle permeabilization [34].

Of note is that adenylylation led to a decrease in the thioflavin T fluorescence in solutions of αSyn incubated under fibrillation conditions (Figure 5). We also observed a change in the fibril morphology of adenylylated αSyn (Figure 6). Typical αSyn fibrils display a bundling pattern that consists of twisted fibrils [49, 50]. We found that adenylyated αSyn fibrils existed largely as bundles of parallel fibrils (Figure 6D), suggesting (as one possibility) that addition of an AMP causes steric hindrance that prevents the fibrils from becoming closely intertwined. Our observation that αSyn adenylylation influences fibril morphology also implies that the difference in thioflavin T fluorescence shown in Figure 5 may reflect a combined effect of, in part, a difference in the quantum yield of the dye bound to the different fibril polymorphs, as well as a decrease in the extent of fibrillation of the adenylylated protein.

Collectively, our findings indicate that αSyn adenylylation interferes with αSyn fibril formation in solution and with αSyn-mediated vesicle disruption, and changes the preferred morphology of the fibrils. Evidence suggests that each of these effects could impact αSyn pathology and interfere with αSyn neurotoxicity [49, 50]. Thus, physiologically, we propose that adenylylation may be triggered as a protective response to cope with the formation of toxic αSyn aggregates.

As mentioned earlier, HYPE plays a key role in regulating the cell’s response to ER stress, and αSyn aggregation is also known to trigger ER stress and the UPR pathway via its interactions with BiP. An important question remains whether HYPE-mediated adenylylation of αSyn occurs in response to ER stress or whether adenylylation of αSyn is independent of UPR activation. Preliminary quantitative RT-PCR analysis of cDNA isolated from human control and PD patient samples revealed that HYPE’s transcriptional levels did not correlate with other UPR associated transcripts (data not shown). Experiments are underway to determine whether HYPE, which we have shown to bind misfolded proteins directly *in vitro*, may bind and adenylylate αSyn independently of the UPR response.

In summary, we have identified αSyn as a novel target for the human Fic protein, HYPE. In doing so, we have uncovered a potential neuroprotective role for Fic-mediated adenylylation in PD that may function concurrently with HYPE’s effects on chaperone activity. It remains to be determined whether the interaction of HYPE with αSyn is connected to ER stress. However, our discovery of a new, physiologically relevant target for HYPE provides significant insights for catalysis and substrate recognition, and opens a new line of research for combatting PD. Given the central role HYPE plays in cell fate, targeting HYPE as opposed to αSyn directly allows a chance to preempt the effects of αSyn aggregation and protect the cell from potential stress-related damage.

## Supplemental Legends

**Figure S1:**
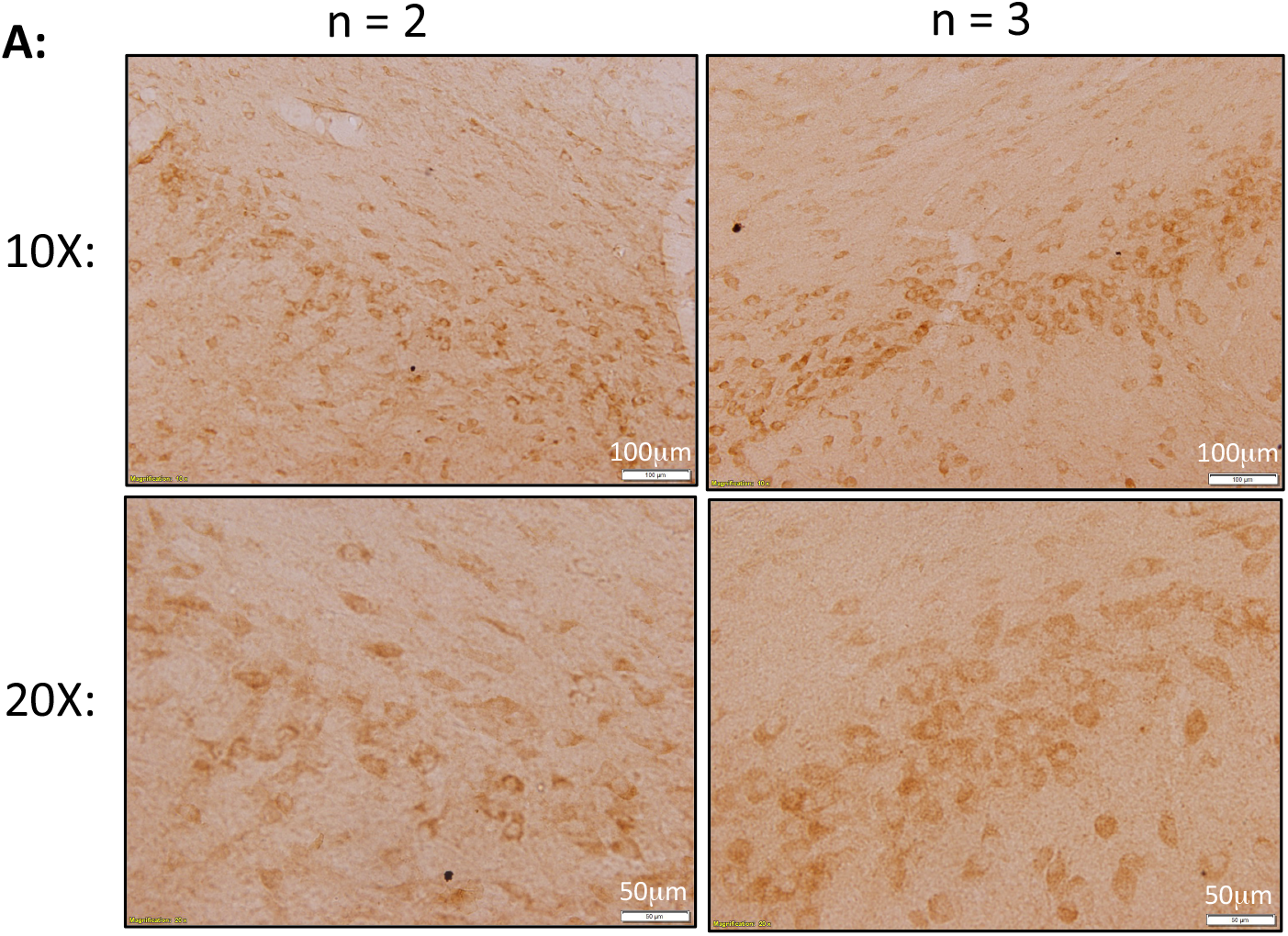
Additional coronal sections of a rat brain (n=2 and n=3) probed with antibody to HYPE and visualized with peroxide-based secondary staining. HYPE-specific staining of the *substantia nigra* is shown at 10X and 20X magnification. Scale bars represent 100 μm and 50 μm, respectively

**Figure S2:**
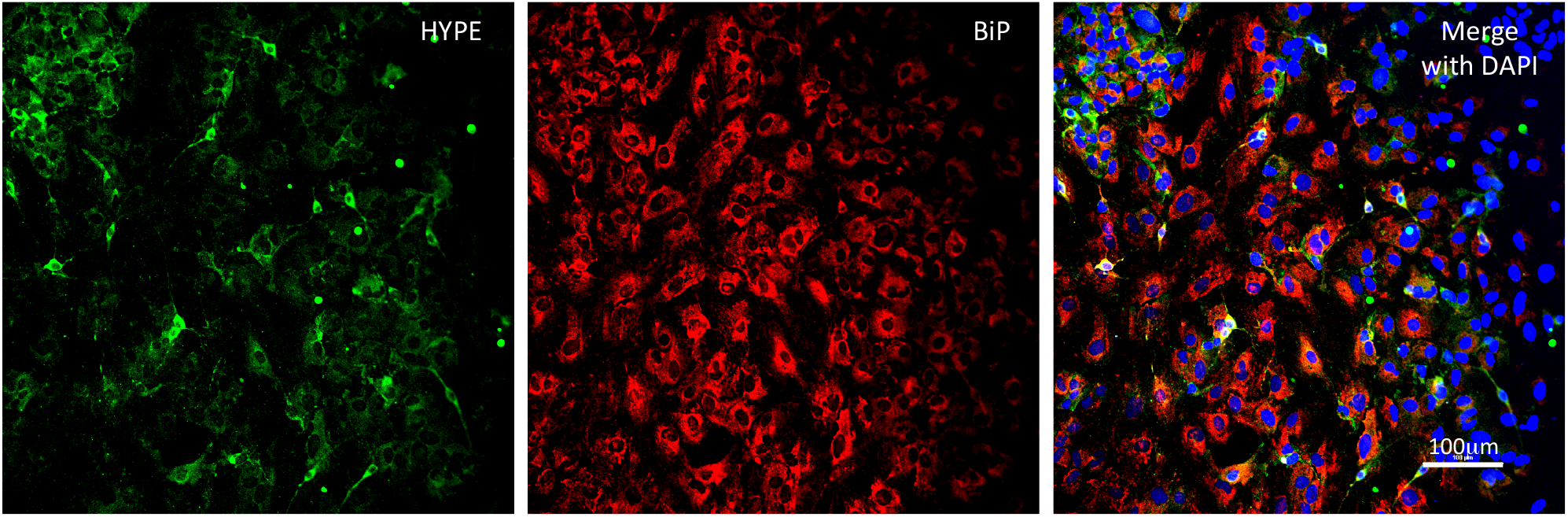
Immunofluorescence images of a rat primary midbrain culture stained for HYPE (green, left panel) or BiP (red, middle panel). The right panel consists of a merged image showing HYPE and BiP immunoreactivity as well as DAPI-stained nuclei. HYPE is preferentially expressed in neurons, whereas BiP staining reflects glial and neuronal expression. Both proteins appear to be preferentially expressed in neuronal cell bodies. Scale bar represents 100 μm.

**Figure S3:**
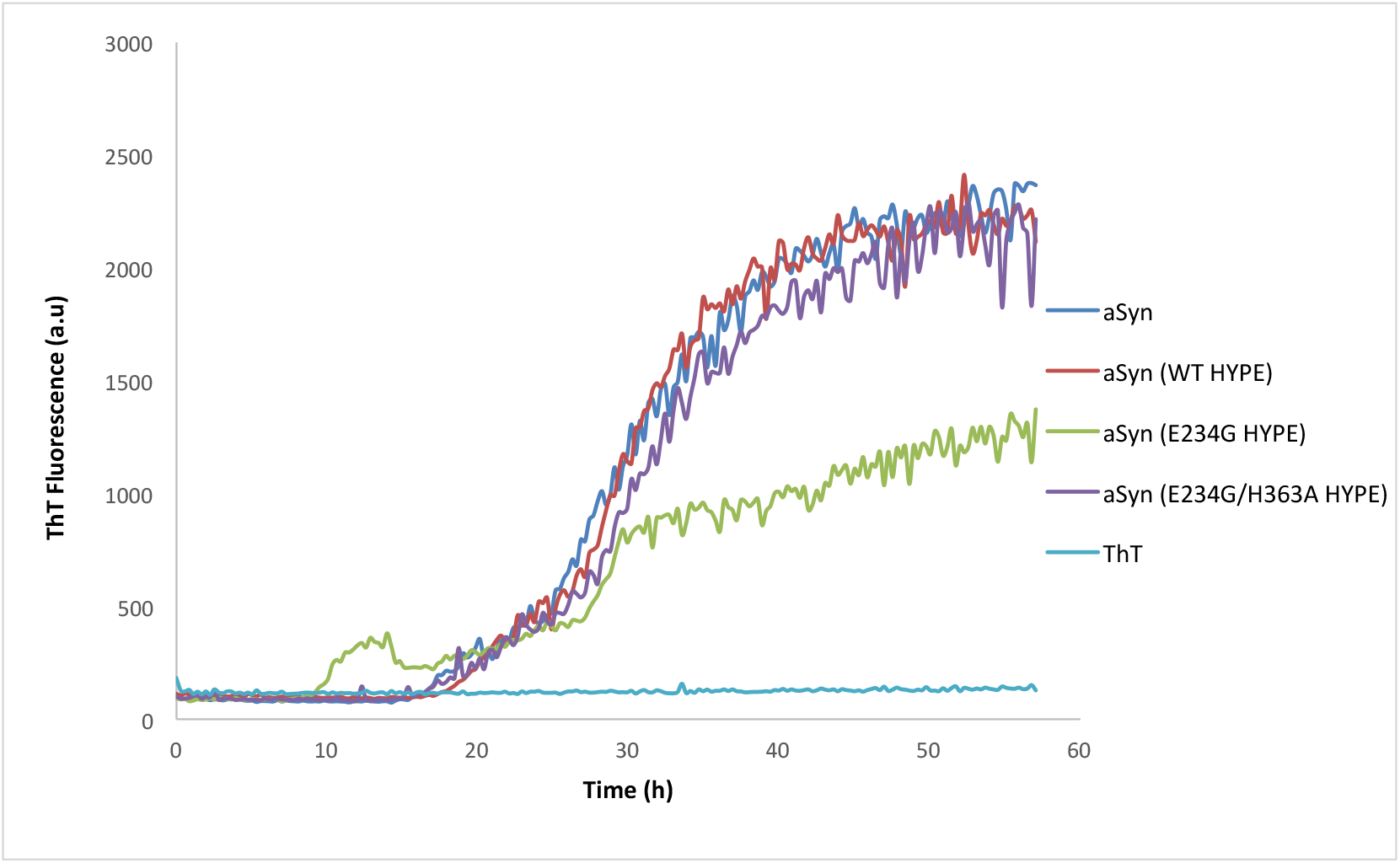
Kinetic traces of a fibrillation assay conducted with purified human WT αSyn adenylylated in the presence of E234G HYPE (green line) or left unmodified as αSyn alone (blue line) show a significant reduction in the yield of ThT fluorescence – potentially reflecting inhibition of fibril elongation – as a result of αSyn adenylylation. Control samples consist of αSyn incubated in the presence of WT-HYPE (red line) or catalytically dead E234G/H363A HYPE (purple line) prior to fibrillation. The background fluorescence for the ThT only control against which all samples were blanked is also shown (light blue line).

## ABBREVIATIONS

AMP: :Adenosine monophosphate
BiP: :Binding immunoglobulin protein
BLI: :Biolayer Inferometry
ER: :Endoplasmic reticulum
FIC: :Filamentation induced by cAMP
GFP: :Green fluorescence protein
GS-AT: :Glutamine synthase adenylyltransferase
HYPE: :Huntingtin yeast interacting protein E
LC-MS: :Liquid chromatography – mass spectrometry
MAP2: :Microtubule associated protein 2
NAC: :Non-amyloid-beta component of Alzheimer’s disease
PD: :Parkinson’s Disease
PTM: :Post-translational modification
SUV: :Small unilamellar vesicles
TEM: :Transmission electron microscopy
TH: :Tyrosine hydroxylase
UPR: :Unfolded protein response
WT: :wild type

## MATERIALS AND METHODS

### Cloning and plasmids

For bacterial expression, WT αSyn/pT7-7 plasmid was procured from Addgene (www.addgene.com; plasmid #36046). This vector was also used as a backbone for cloning His_6_-tagged WT-αSyn by PCR amplification.

### Protein expression and purification

Untagged human WT αSyn, its A53T variant, and mouse WT αSyn purified as previously described [34]. Oligomeric species of untagged αSyn were separated from low molecular weight, mostly monomeric, species by passing the protein over a 100kDa cut-off filter. His_6_-tagged αSyn was expressed in *E. coli* BL21-DE3 (Stratagene) in LB medium containing 100 μg/ml of ampicillin to a density of A_600_ = 0.6. Protein expression was induced for 3 hours at 37°C with 0.4 mM IPTG. Cells were lysed in lysis buffer (50 mM HEPES pH 8.0, 250 mM NaCl, 5 mM imidazole, 10% (v/v) glycerol) containing 1mM PMSF and purified using a cobalt resin. Resin was washed with wash buffer (50 mM HEPES pH 8.0, 250 mM NaCl, 40 mM imidazole,10% (v/v) glycerol) and αSyn-His protein was eluted from the resin using elution buffer (50 mM HEPES pH 8.0, 250 mM NaCl, 350 mM imidazole, 10% (v/v) glycerol) and further purified by ion exchange chromatography using 100 mM Tris-HCl pH 7.5 with a salt gradient from 10 mM-1M NaCl. Fractions containing αSyn were purified by size exclusion chromatography in buffer containing 100 mM Tris-HCl pH 7.5, 100 mM NaCl to separate monomer and the void-volume oligomer. Protein concentrations were measured using the Bradford method. Purity was determined by SDS-PAGE and proteins were stored at −80°C.

Recombinant HYPE and its mutants were purified as previously described [7]. Specifically, wild type and HYPE mutant proteins were expressed in *E. coli* BL21-DE3-RILP (Stratagene) in LB medium containing 50 μg/ml of kanamycin (pSMT3) to a density of A_600_ = 0.6. Protein expression was induced for 12-16 hours at 18°C with 0.4 mM IPTG. Cells were lysed in lysis buffer (50 mM HEPES pH 8.0, 250 mM NaCl, 5 mM imidazole) containing 1 mM PMSF protease inhibitor and purified using a cobalt resin. Resin was washed with wash buffer (50 mM HEPES pH 8.0, 250 mM NaCl, 20 mM imidazole). His_6_-SUMO tag was cleaved by incubating proteins with ULP1 at 4°C. The protein mixture was diluted in wash buffer without imidazole and re-applied to a cobalt column, and flow through containing HYPE was further purified by ion exchange chromatography using 100 mM Tris-HCl pH 7.5 with a salt gradient from 10 mM – 1 M NaCl. Fractions containing HYPE were purified by size exclusion chromatography in buffer containing 100 mM Tris-HCl pH 7.5, 100 mM NaCl. Protein concentrations were measured using the Bradford method. Purity was determined by SDS-PAGE and proteins were stored at −80°C.

### Biolayer interferometry

The binding kinetic assays were performed on an Octet Red 384 instrument (ForteBio, Menlo Park, CA) with the Biolayer Interferometry (BLI) technique. The assays were carried out using Anti-pentaHis (HIS1K) biosensors (ForteBio, Menlo Park, CA) at 30 °C with the plate shaking speed at 1000 rpm. HIS1K sensors were dipped into the wells of 20 mM His_6_-αSyn in HEPES buffer (50 mM HEPES, 150 mM NaCl, pH7.5) for 2 minutes to load αSyn on the sensors, followed by incubation in buffer for a minute to build a baseline. Next, the sensors were dipped into wells of N-terminally His tagged HYPE in HEPES Buffer at a concentration ranging from 1 μM to 150 μM for 2 minutes to obtain association curves. The sensors were then dipped back to the baseline HEPES buffer wells for 3 minutes to obtain dissociation curves. The data were analyzed by the ForteBio data analysis software v.9.0.0.12. The binding constants (Kd) and kinetic parameters were determined by globally fitting data to the 1:1 fitting model.

### *In vitro* adenylylation assays

0.5 μg of E234G or E234G/H363A HYPE (aa 102-458) protein was incubated either alone or with 5 μg of untagged αSyn monomer or oligomer in an adenylylation reaction containing 5 mM HEPES pH 7.5, 1 mM manganese chloride tetrahydrate, 0.5 mM EDTA and 0.01 mCi α-^32^P-ATP for 15 minutes. Reactions were stopped with SDS PAGE loading buffer, separated on 4-20% polyacrylamide gels and visualized by autoradiography.

### Mammalian cell culture assays

HEK293T cells were cultured in Dulbecco’s Modified Eagle Medium (Sigma-Aldrich) supplemented with 5% (v/v) fetal bovine serum (Life Technology) and 100 g/ml penicillin/streptomycin (Sigma-Aldrich) at 37°C with 5% CO2. Fugene HD (Promega) was used for transfections according to the manufacturer’s directions. Cells were transfected for 30 hrs. Primary midbrain cultures were prepared and treated as previously described [34].

### Mass spectrometry

For in-gel protein digestion, gel bands were cut out, destained, reduced with 5mM DTT and alkylated with 10mM iodoacetamide and digested with either trypsin at 37C or chymotrypsin at room temperature essentially as described by Shevchenko *et al*. [51] with minor modifications. For LC-MS/MS analysis, the dried peptide mix was reconstituted in a solution of 2% formic acid (FA) for MS analysis. Peptides were loaded with the autosampler directly onto a 2cm C18 PepMap pre-column (Thermo Scientific, San Jose, CA) which was attached to a 50cm EASY-Spray C18 column (Thermo Scientific). Peptides were eluted from the column using a Dionex Ultimate 3000 Nano LC system with a 1.5 min gradient from 2% buffer B to 5% buffer B (100% acetonitrile, 0.1% formic acid), followed by a 15 min gradient to 20% of B, followed by a 2 min gradient to 30% of B. The gradient was switched from 30% to 85% buffer B over 1 min and held constant for 1 min. Finally, the gradient was changed from 85% buffer B to 98% buffer A (100% water, 0.1% formic acid) over 1 min, and then held constant at 98%t buffer A for 15 more minutes. The application of a 2.0 kV distal voltage electrosprayed the eluting peptides directly into the Thermo Fusion Tribrid mass spectrometer equipped with an EASY-Spray source (Thermo Scientific). Mass spectrometer-scanning functions and HPLC gradients were controlled by the Xcalibur data system (Thermo Scientific). Initial experiments were performed to identify AMP modified peptides by running each digest separately with a data dependent method to acquire as many MS/MS in a 3 second span. This data was analyzed for the presence of AMP modification with either inspect (proteomics.ucsd.edu) or Thermo Proteome Discoverer 2.1 and looking for a modification of 329.05252 on either tyrosine, threonine or serine or by manually looking at MS/MS scans and looking for the specific signal for AMP (m/z 348.08). Three MS/MS scans were identified that contained AMP, however, the most abundant peak was identified as the neutral loss of AMP from the parent mass. Therefore an analysis was conducted that targeted these neutral loss ions for an additional MS3 scan to identify the peptide sequence and position of the AMP modification.

### Immunocytochemistry and immunohistochemistry (IHC)

Primary midbrain cultures were fixed with 4% (w/v) paraformaldehyde and blocked with 1X PBS containing 10% (v/v) Fetal Bovine Serum (FBS), 1% (w/v) BSA and 0.3% (v/v) Triton X-100 for 1 hour. This was followed by treatment with primary antibodies - mouse anti-HYPE (Abcam cat# ab67163, 1:100), chicken anti-MAP2 (Encor cat# CPCA-MAP2, 1:2000) or rabbit anti-TH (Phosphosolutions cat# 2025-THRAB, 1:1000) diluted in 1X PBS containing 1% (w/v) BSA and incubated overnight at 4°C. The cells were washed and incubated with secondary antibodies Alexa anti-chicken 594, Alexa anti-rabbit 488 or Alexa anti-mouse 594 for 1 hour at room temperature, washed, and imaged using the CYTATION 3 Cell Imaging Reader or Nikon Ti confocal microscope

Rat brain sections (40 μm) were obtained using a microtome and stored in cryo-protectant solution at −20°C. On the day of the experiment, sections were transferred to a separate plate and washed six times with PBS. All incubations were conducted while shaking at 100 rpm. The sections were blocked with 10% Normal Donkey Serum (NDS) in 0.3% PBS-Triton for 90 minutes, treated with mouse primary antibody for HYPE (Abcam cat# ab67163, 1:100) diluted in PBS containing 1% NDS and 0.3% Triton for 24 hours at 4°C. The following day, the sections were washed three times with PBS, incubated with biotinylated secondary antibody anti-mouse (Jackson ImmunoResearch, cat# 715-065-151, 1:200) for 90 minutes at room temperature, washed with PBS and incubated in the premade ABC solution from VECTASTAIN Elite ABC HRP Kit (Vector Laboratories, cat# PK-7200) for 60 minutes. A solution of DAB (diaminobenzidine) and H_2_O_2_ was prepared using the DAB Peroxidase Substrate Kit (Vector Laboratories, cat# SK-4100). The sections were were then treated with the DAB solution until a brown color developed, washed to remove non-specific staining and transferred to a slide. The tissues were allowed to dry overnight and were dehydrated using a series of solutions of increasing ethanol concentrations (70-100%), followed by treatment with Histoclear, mounted using Histomount and dried overnight before microscopy using the Olympus BX53 light microscope.

### αSyn fibrillation

Lyophilized untagged αSyn was dissolved in PBS (pH 7.4) with 0.02% [w/v] NaN_3_, and dialyzed against the same buffer (24 h at 4°C) to remove excess salt. The solution was filtered by successive centrifugation steps through a 0.22 μm spin filter and a 100 kDa centrifugal filter to isolate low molecular weight (mostly monomeric) protein. This low molecular weight αSyn was incubated in PBS for an hour with or without HYPE as outlined above. Fibrillation solutions of αSyn were prepared by diluting the adenylation reaction mixture (or blank) to a final concentration of 35 μM in the wells of a 96-well plate. To determine the extent of αSyn fibrillation, thioflavin T (ThioT, final concentration, 20 μM) was added to the fibrillation solutions, which were incubated at 37 °C with shaking at the ‘normal’ setting in a Spectra Fluor Plus or Genios plate reader (Tecan, Upsala, Sweden). HYPE was not removed from the fibrillation mixture as it did not interfere with fibrillation or show increased ThioT fluorescence on its own. A nylon ball was added to avoid inhomogeneous mixing of samples. ThioT fluorescence was measured with excitation at 440 nm and emission at 490 nm. Mean ThioT fluorescence data determined from 2-4 technical replicates were normalized using the following equation:

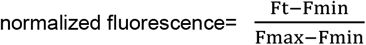

where F_t_ is the ThioT fluorescence emission intensity at time t, and F_min_ and F_max_ are the minimum and maximum fluorescence intensities during the incubation, respectively.

### Synthetic vesicle disruption assay and on-bead adenylylation

Calcein loaded Small unilamellar vesicles (SUVs) (diameter ~ 90 nm) were prepared as described previously [37]. We opted to examine SUVs composed of egg PG and egg PC (1:1 mol/mol) because they contain anionic lipids necessary for αSyn membrane interactions and this lipid composition is compatible with producing stable 90 nm vesicles. Egg PG and egg PC suspended in chloroform were mixed in a round bottom flask. The chloroform was evaporated under a nitrogen stream and further dried under vacuum for 1 h. Dried lipids were suspended in a solution of calcein (170 mM) prepared in water using NaOH to adjust the pH to 7.4. The final osmolality was 280 mOsm/kg. The liposomes were sized by extruding the suspension through a 50 nm pore size Whatman membrane. Calcein-containing vesicles were isolated from free calcein via gel filtration through a Sephadex G-50 column pre-equilibrated with PBS, pH 7.4, 0.02 % (w/v) NaN3 (280 mOsm/kg). Fractions containing isolated vesicles were pooled and stored at 4 °C until use. Calcein-loaded vesicles were found to be very stable, with no spontaneous dye leakage observed over several weeks. The size of the vesicles was confirmed by dynamic light scattering.

To avoid any interference of HYPE on vesicle permeabilization, the adenylation reaction was done in PBS for one hour as mentioned previously (the adenylation method need to added at the beginning) but with Ni-agarose bead bound HYPE. Untagged αSyn was removed from the beads by removing the supernatant after centrifugation of the samples. To remove any non-specifically bound αSyn from the beads, the sample was washed with small volumes of PBS and pooled with the supernatant as mentioned above. αSyn concentration was calculated with BCA assay and absorbance measurement at 280nm.

For membrane disruption experiments, monomeric αSyn variants (40 μM, unmodified and modified) were incubated with calcein-loaded PG:PC SUVs (protein-to-lipid ratio, 1:20 mol/mol) at 37°C in a total volume of 40 μL. Samples were analyzed with a TECAN fluorescence plate reader (excitation wavelength, 485 nm; emission wavelengths, 530 nm). To determine the maximum dye release, vesicles were lysed by adding 3 μL of 1 % (v/v) Triton X-100. The percentage leakage at time t was determined using the following equation:

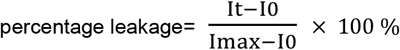

where It is the fluorescence emission intensity at time t, I_0_ is the intensity at time 0, and Imax is the maximal intensity determined after detergent lysis of the vesicles. αSyn-mediated leakage reached a plateau after 12-16h, and the plateau value was used to compare the leakage among the HYPE-modified and unmodified samples.

### Western blot analysis of fibril samples

A 150 L aliquot of the fibrillation sample was centrifuged for 60min at 100,000 x g (Beckman Coulter Optima MAX-XP) to precipitate the fibrils. The supernatant was collected to analyze the residual monomeric content. The fibril pellet was dissolved in 150 L PBS and vortexed gently and used for analysis. Appropriate amounts of the samples were mixed with Laemmli sample buffer and denatured proteins were separated via SDS-PAGE on a 12 %(w/v) polyacrylamide gel and transferred to an Immobilon-FL PVDF membrane (pore size, 0.45 m). The membrane was probed with mouse anti-αSyn (BD Biosciences; 1:1,500), and an antimouse AP-linked secondary antibody (Cell Signaling Technology, 1: 3,000). To visualize the bands, the membrane was exposed to ECF substrate and imaged using a Typhoon imaging system (GE Life Sciences, Pittsburgh, PA). Band intensities were quantified using ImageJ software (NIH, Bethesda, MD), and averages calculated by quantifying each band in triplicate.

### Transmission Electron Microscopy (TEM) analysis of αSyn amyloid-like fibrils

The morphology of the αSyn amyloid-like fibrils was analyzed by negative stain biological TEM. 3 L of the αSyn sample solution of interest (35 M, directly from the fibrillation assay) was pipetted on a discharged carbon-coated copper TEM grid (Electron Microscopy Sciences, CF400CU; 400 mesh) and incubated for 1 min. Subsequently, the sample was washed with deionized water carefully without letting it dry and stained with a 1% (w/v) phosphotungstic acid (PTA) solution (3 L), which was left in contact with the protein on the grid for 1 min. The excess solution was then removed by blotting with filter paper, and the sample was imaged using an FEI Tecnai G2 20 Transmission Electron Microscope operating at 200 KV.

## Acknowledgments

We thank the Biophysical Analysis Laboratory at Purdue University’s Bindley Biosciences Center for use its Octet Red 384 for BLI analysis, and Dr. Chris Gilpin at the Purdue Electron Microscopy Core for use of the FEI Tecnai G2 20 TEM. Finally, we are grateful to members of the Mattoo and Rochet labs for their helpful discussions.

## Author contributions

S.M. conceived the study. S.M., J-C.R., A.S., and S.D. designed the study. A.S., S.D., A.C., A.C., and B.G.W. conducted the experiments. D.Y. and P.M. provided initial samples of purified αSyn and followed protocols developed by J.R.C. for handling rat brain samples. R.S. helped with TEM experiments. A.K. conducted mass spectrometric analyses. S.M. and J-C.R. supervised the study. S.M. and A.S. wrote the manuscript. All authors revised and agreed to the final version of the manuscript.

